# Integrating single-cell RNA-seq and imaging with SCOPE-seq2

**DOI:** 10.1101/2020.06.28.176404

**Authors:** Zhouzerui Liu, Jinzhou Yuan, Anna Lasorella, Antonio Iavarone, Jeffrey N. Bruce, Peter Canoll, Peter A. Sims

## Abstract

Live cell imaging allows direct observation and monitoring of phenotypes that are difficult to infer from transcriptomics. However, existing methods for linking microscopy and single-cell RNA-seq (scRNA-seq) have limited scalability. Here, we describe an upgraded version of Single Cell Optical Phenotyping and Expression (SCOPE-seq2) for combining single-cell imaging and expression profiling, with substantial improvements in throughput, molecular capture efficiency, linking accuracy, and compatibility with standard microscopy instrumentation. We introduce improved optically decodable mRNA capture beads and implement a more scalable and simplified optical decoding process. We demonstrate the utility of SCOPE-seq2 for fluorescence, morphological, and expression profiling of individual primary cells from a human glioblastoma (GBM) surgical sample, revealing relationships between simple imaging features and cellular identity, particularly among malignantly transformed tumor cells.

## Introduction

High-throughput single-cell RNA-sequencing (scRNA-seq) has revolutionized molecular profiling of complex tissues and cell state transitions like differentiation (Bose et al., 2015; Klein et al., 2015; Macosko et al., 2015). However, many cellular phenotypes are difficult to infer from the transcriptome. Furthermore, scRNA-seq is fundamentally an end-point measurement, and does not enable real-time monitoring of individual cells. However, cellular imaging by microscopy can be applied to live cells for direct measurement and monitoring of numerous cellular phenotypes such as protein abundance and localization, cellular morphology, electrical activity, active transport, and enzymatic and metabolic activity, taking advantage of a vast number of fluorescent probes that have been developed over decades. Therefore, the ability to link cellular phenotypes measured by microscopy directly to the gene expression profiles of individual cells would allow a more comprehensive description of cellular states (Lane et al., 2017; Yuan et al., 2018b).

Previous methods that link optical measurements with scRNA-seq have technical limitations. Early scRNA-seq methods used fluorescence activate cell sorting (FACS) to deposit cells into individual wells of a standard multi-well plate and prepare cDNA libraries from each captured cell one at-a-time (Hochgerner et al., 2017; Lane et al., 2017; Shalek et al., 2013). While these methods can link cytometric and even imaging data to scRNA-seq, they lack scalability. Recent advances have combined microfluidics and barcoded mRNA capture beads to facilitate pooled library preparation from thousands of individual cells, which reduces costs and increases scalability, but these methods lack the ability to link cellular images to sequencing data (Bose et al., 2015; Klein et al., 2015; Macosko et al., 2015). Single Cell Optical Phenotyping and Expression (SCOPE-seq) is a scalable method for linking scRNA-seq with live cell imaging in which individual cells are captured in microwells, imaged, and then co-encapsulated with barcoded mRNA capture beads for pooled scRNA-seq (Yuan et al., 2018b). In addition to the cell-identifying barcode that is incorporated into the cDNA of each cell, the SCOPE-seq beads included a second barcode sequence that could be optically decoded by cyclic hybridization of fluorescently-labeled oligonucleotide probes. While successful, this approach had limited throughput, mRNA capture efficiency, and required a complex procedure for linking the two barcodes.

Here, we present SCOPE-seq2, which significantly improves the throughput, mRNA capture efficiency, and accuracy of the previously published SCOPE-seq technology(Yuan et al., 2018b) and uses a simpler and more scalable approach for optical decoding. We thoroughly characterize the performance of SCOPE-seq2, which enables applications in primary cells dissociated from tissues. Finally, we demonstrate SCOPE-seq2 profiling of a human glioblastoma (GBM) surgical specimen and identify relationships between basic imaging features and the lineage identities of transformed tumor cells.

## Results

### SCOPE-seq2 Methodology

After dissociating cell cultures or tissues into single cell suspensions and staining the cells with selected fluorescent dyes, we processed cells using the SCOPE-seq2 workflow (Figure 1A). SCOPE-seq2 consists of five steps: (1) capture individual cells in microfabricated microwells by limiting dilution; (2) measure the desired phenotypes of each cell by imaging; (3) capture mRNA and synthesize barcoded cDNA from each cell on an improved optically decodable bead; (4) optically decode cell barcode sequences for linking imaging and sequencing by sequential probe hybridization; (5) amplify and sequence barcoded cDNA to obtain an expression profile for each cell (Figure 1A).

**Figure 1.**
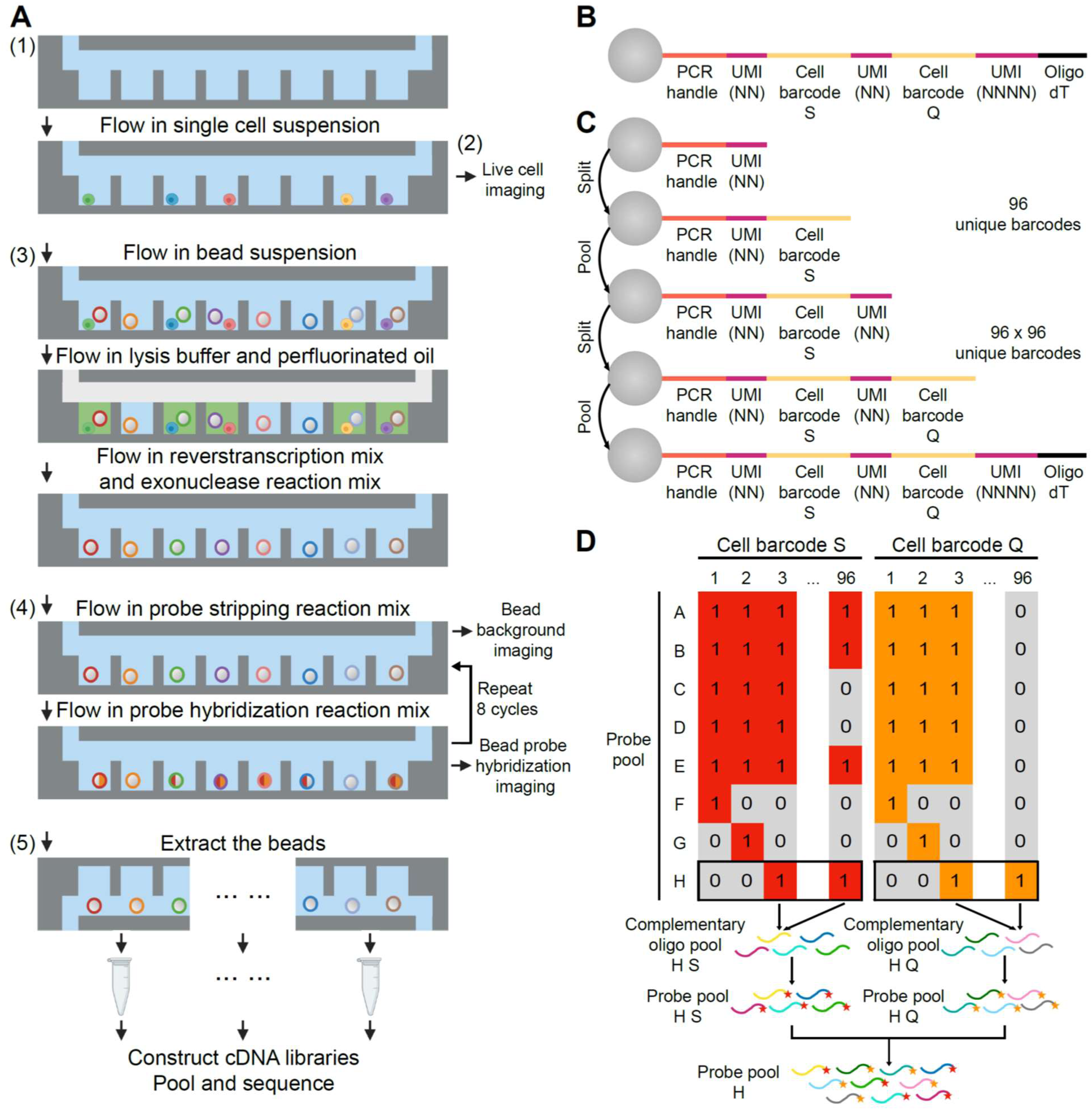
Overview of SCOPE-seq2. A) A schematic representation of the experimental workflow for SCOPE-seq2. B) Oligonucleotide design for SCOPE-seq2 optically decodable mRNA capture beads. C) Split-pool synthesis scheme for generating combinatorial SCOPE-seq2 barcodes with the structure shown in B). D) Schematic for generating pools of fluorescent probes for SCOPE-seq2 optical decoding.

In conventional scRNA-seq, the barcoded mRNA capture beads are randomly co-encapsulated with individual cells in the microwells, so we do not know which barcode is incorporated into the cDNA library of each imaged cell. However, in SCOPE-seq we can identify the barcode sequence on the bead in each microwell by hybridizing fluorescently-labeled oligonucleotide probes and imaging the beads with a fluorescence microscope (optical decoding). In our original report, the cell-identifying barcode that was incorporated into the cDNA library of each cell and the optically decodable barcode sequence were distinct, and we had to prepare a separate sequencing library to link the two sequences. In addition, only a small subset of oligonucleotides on each bead actually contained the optically decodable barcode sequence, limiting the fluorescence signal and therefore imaging speed and throughput. For SCOPE-seq2, we devised an improved optically decodable bead where the sequencing and optically decodable barcode sequences that identify a given cell are the same (Figure 1B). The cell barcode contains two 8-nucleotide sequences, each of which is a member of a pool of 96 sequences (Table EV 1). An 8-nucleotide random sequence is dispersed into three parts and serves as both a unique molecular identifier (UMI) and a spacer between other functional sequences on the bead. The oligonucleotides on all beads share two common sequences - a universal PCR adapter on the 5’-end and oligo(dT) on the 3’-end for mRNA capture and cDNA amplification. The oligonucleotides are synthesized by split-pool, solid-phase synthesis (Figure 1C). Beads are pooled together to add common sequences and random UMIs, and are split into 96 reactions to add one of the 96 cell barcode sequences. After two rounds of split-pooling, a total of 96^2^ = 9,216 cell barcodes are generated. To generate cDNA from cells, we co-encapsulate the cells with these beads, lyse the cells, capture cell mRNAs on beads by hybridization, and reverse transcribe the captured mRNAs.

To link cellular imaging with scRNA-seq from the same cell, we identify the cell barcode sequence on each bead in the microwell array by sequential fluorescent probe hybridization. Our strategy is related to methods of decoding DNA microarrays and highly multiplexed fluorescence *in situ* hybridization (FISH) (Chen et al., 2015; Gunderson et al., 2004; Lubeck et al., 2014; Shah et al., 2016). We use a temporal barcoding strategy in which each 8-nt cell barcode sequence corresponds to a unique, pre-defined 8-bit binary code (Table EV2, Table EV3). Each bit of the binary code can be read out by one cycle of probe hybridization, where the presence or absence of a hybridized probe indicates one or zero, respectively. The two parts of the cell barcode can be decoded simultaneously using two sets of differently colored fluorescent probes. To realize this decoding scheme, we generate a pool of fluorescent probes for each cycle of hybridization. All oligos whose sequences are complimentary to the cell barcode sequence marked ‘1’ in the corresponding binary code are pooled and conjugated with fluorophores, Cy5 or Cy3. Distinct fluorophore-conjugated probes against the two 8-nucleotide sequences comprising the cell barcode are then pooled together to form the final probe pool (Figure 1D). Thus, we are able to decode all possible cell barcode sequences by eight cycles of two-color probe hybridization. This approach is more scalable than the original SCOPE-seq strategy and gives a brighter signal on the bead surface because every primer contains an optically decodable barcode. Thus, SCOPE-seq2 beads are compatible with higher speed imaging, leading to higher throughput.

Finally, we further increased the cell indexing capacity to 96^2^ × 10 = 92,160 by dividing the microwells into ten regions as previously described (Yuan et al., 2018b). We extract the beads from each region of the device separately for library construction and indexing, and then sequence the cDNA libraries from each region in a single pool.

### Cell barcode optical decoding analysis for SCOPE-seq2

To decode the cell barcode sequences from imaging, a ‘cycle-by-cycle’ method was used in SCOPE-seq (Gunderson et al., 2004; Yuan et al., 2018b), which calls the binary code for each bead based on the bimodal distribution of intensity values across all beads in each hybridization cycle. This method works well when the bead fluorescence intensity values of the ‘one’ state population are well separated from that of the ‘zero’ state population. However, because the beads exhibit auto-fluorescence at shorter wavelengths, the two populations are not as well separated in the Cy3 emission channel as in the Cy5 emission channel (Figure EV1).

To accurately decode the cell barcode sequences from imaging, we utilized a modified ‘bead-by-bead’ fluorescence intensity analysis strategy, which has been used to decode randomly ordered DNA microarrays (Gunderson et al., 2004). We determine the cell barcode sequences of each bead by sorting the eight intensity values in each emission channel in ascending order, calculating the relative intensity change between each pair of adjacent values, establishing a threshold based on the largest relative intensity change to assign a binary code, and mapping the binary code to the actual cell barcode sequence (Figure 2A). For those unmappable binary codes, we repeatedly re-assign the binary code based on the next largest relative intensity change until the code can be successfully mapped to a cell barcode sequence. Since this method decodes each bead independently, we expected that it would give better results when the ‘one’ and ‘zero’ intensity states were poorly separated.

**Figure 2.**
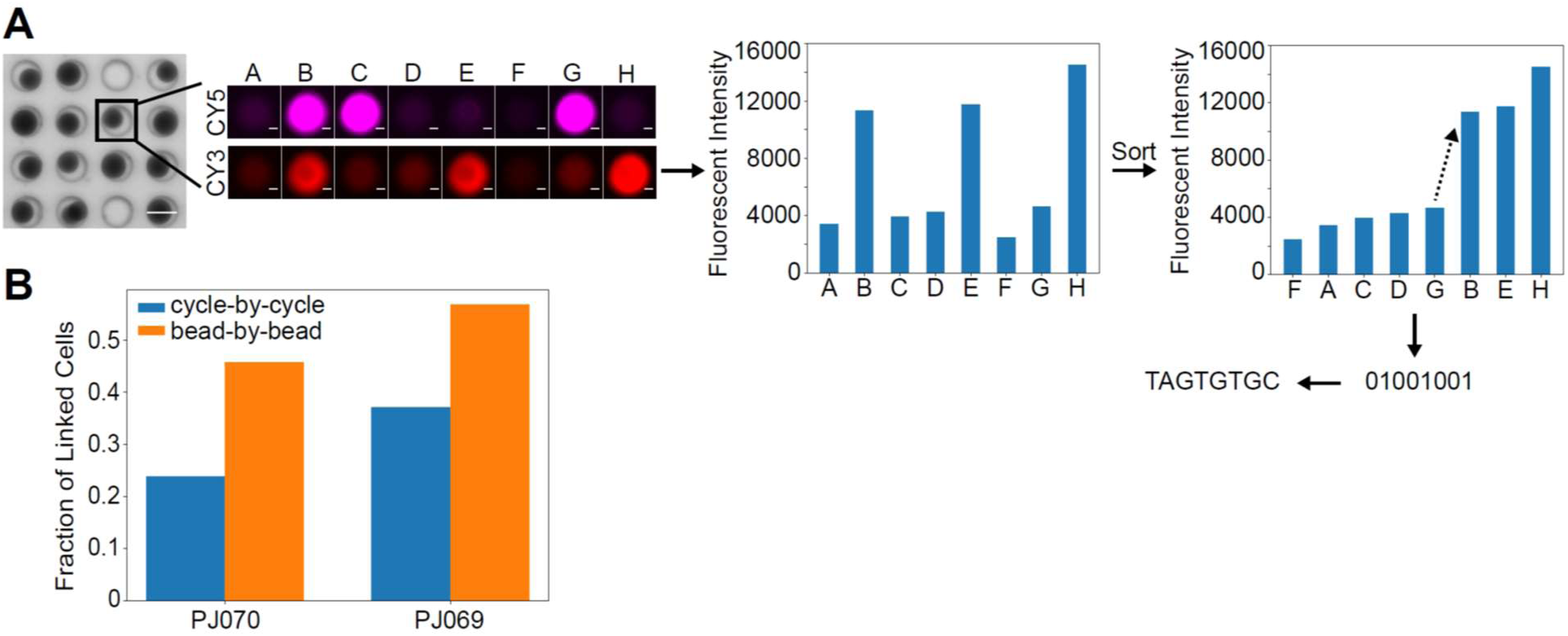
Optical decoding of cell barcodes. A) Bright field image of SCOPE-seq2 beads in PDMS microwells (left) and two-color fluorescence images of a SCOPE-seq2 bead after each cycle of optical decoding (right). Scale bars: 50 μm (multi-well image, left) and 10 μm (single-well images, right). Bar plots show the 8-cycle fluorescent intensity values before (left) and after sort (right) of a SCOPE-seq2 bead in the CY3 emission channel. An arrow shows the two adjacent values with the largest relative intensity change. B) Comparison of the ‘bead-by-bead’ and ‘cycle-by-cycle’ decoding methods. A bar plot shows the fraction of scRNA-seq expression profiles that are successfully linked to cell images in two different experiments (PJ069 and PJ070).

In SCOPE-seq2, we only used 96 out of 2^8^ = 256 possible binary codes, and more importantly, the number of sequenced cell barcodes (< 10,000 cells per experiment) is much fewer than the total 92,160 possible cell barcodes. Therefore, an error in optical decoding would mainly result in assigning the bead an unmappable binary code, or a cell barcode that does not appear in the sequencing data. Both kinds of misassignments further lead to the failure of linking imaging and sequencing data sets rather than incorrect linking. Thus, a more accurate optical decoding method would give a higher fraction of linked imaging and sequencing data. To compare the ‘bead-by-bead’ method with the ‘cycle-by-cycle’ method, we tested these two methods on two datasets. In dataset PJ070 and PJ069, we linked 46% and 57% scRNA-seq profiles with cell images using the ‘bead-by-bead’ method compared to only 24% and 37% using the ‘cycle-by-cycle’ method. In both datasets, we observed at least a 20% increase in the fraction of linked cells with the ‘bead-by-bead’ method (Figure 2B), which suggests that the ‘bead-by-bead’ method is more suitable for cell barcode optical decoding in SCOPE-seq2.

### Validation of SCOPE-seq2

To demonstrate the performance of SCOPE-seq2, in terms of throughput, molecular capture efficiency, and accuracy of linking imaging and sequencing data, we performed an experiment with mixed human (U87) and mouse (3T3) cells labeled with two differently colored live staining dyes. We loaded the mixed cells into the microwells at a relatively high density and obtained 9,061 transcriptional profiles from a single experiment. At saturating sequencing depth, we detected on average 10,245 mRNA transcripts from 3,548 genes per cell (Figure 3A, 3B), which is a nearly three-fold improvement over the original report of SCOPE-seq (Yuan et al., 2018b). To evaluate the linking accuracy of SCOPE-seq2, we identified the species of each cell from the color of the fluorescent label and from the species-specific alignment rate in RNA-seq (a cell with >90% of reads aligning to the transcriptome of a given species was considered species-specific), and examined the consistency of the two cell species calls. In the 4,145 scRNA-seq profiles that we successfully linked with imaging data, we obtained a class-balanced linking accuracy of 99.2% (0.8% error rate), with 98.8% of human cells and 99.6% of mouse cells agreeing with the species calls from two-color imaging (Figure 3C). This represents a nearly five-fold improvement in accuracy over the original report of SCOPE-seq (3.9% error rate). We are also able to confidently remove multiplets in SCOPE-seq2 by manually identifying mixed-species and single-species multiplets from the two-color cell images (Figure EV2). By comparing image-based and sequencing-based mixed-species multiplets, we obtained a multiplet detection sensitivity of 68.8% and a specificity of 97.0%. A large portion of transcriptional profiles with low purity have been removed (Figure 3D). Since we confirmed that SCOPE-seq2 has high linking accuracy, we suspected that the mixed-species multiplets detected by sequencing but not imaging were because of the imperfections in scRNA-seq data, which served as our ground truth.

**Figure 3.**
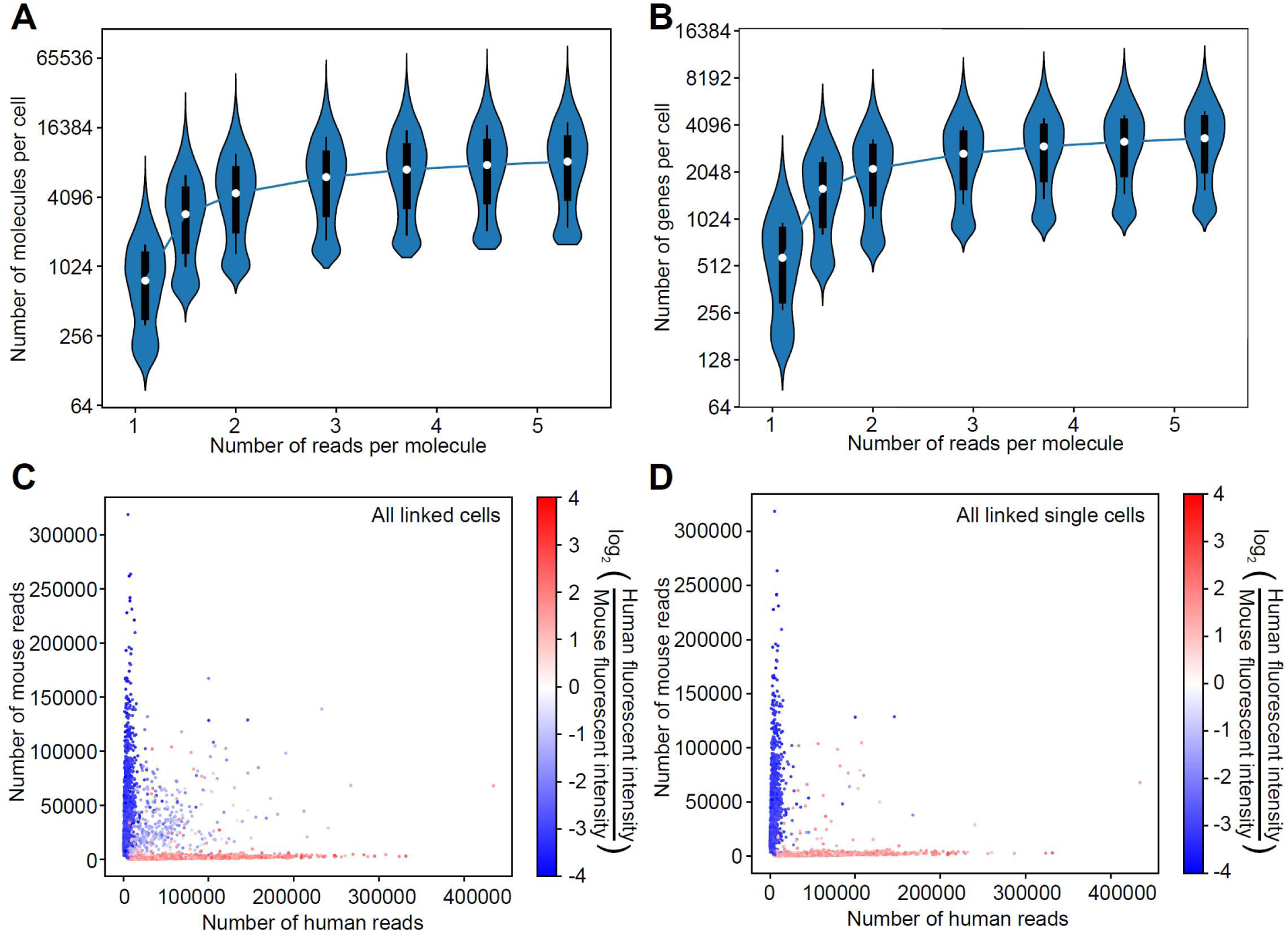
Validation and performance of SCOPE-seq2 in a mixed-species experiment. Saturation analysis of the number of A) unique transcript molecules and B) genes detected per cell (violin plots indicate distributions across cells). Scatter plots showing the number of uniquely aligned human and mouse reads corresponding to each cell barcode linked to images, before C) and after D) removal of multiplets. Each point (cell) is colored by the fluorescence intensity ratio of the human and mouse live staining channels, indicating excellent agreement between scRNA-seq and imaging.

### SCOPE-seq2 allows paired analysis of image-based and transcriptional phenotypes in individual cells isolated from human tissues

To demonstrate that we can collect paired optical and transcriptional phenotypes from human tissue samples using SCOPE-seq2, we performed an experiment on cells dissociated from a human GBM surgical sample and labeled with calcein AM, a fluorgenic dye that reports esterase activity. We obtained 1,954 scRNA-seq profiles and linked 1,110 of them to live cell images. We manually removed cell multiplets based on imaging. Calcein AM is commonly used as a live stain, and so we also removed outlier cells with low fluorescence intensity (see Methods). Malignantly transformed GBM cells often resemble non-neoplastic neural cell types in the adult brain, and so simple marker-based analysis is insufficient to confirm malignant status. To address this, we identified a large population of cells with amplification of chromosome 7 and loss of chromosome 10, two commonly co-occurring aneuploidies that are pervasive in GBM (Phillips et al., 2006; Yuan et al., 2018a), based on the gene expression (Figure EV3). We then computed a low-dimensional representation of the data using single-cell hierarchical Poisson factorization (scHPF) to identify key gene signatures that define the population (Levitin et al., 2019) and visualized their distributions across cells using Uniform Manifold Approximation and Projection (UMAP). We recovered all of the major cell types that have been previously reported from scRNA-seq of GBM (Neftel et al., 2019; Yuan et al., 2018a) including myeloid cells, endothelial cells, pericytes, malignant-transformed astrocyte-like cells, mesenchymal-like cells, oligodendrocyte-progenitor-like/neuroblast-progenitor-like cells (OPC/NPC) and cycling cells (Figure 4A, 4B). We also measured 16 imaging features from cell images and grouped those features into three categories, cell size, shape and calcein AM intensity using unsupervised hierarchical clustering (Figure 4C) to create three imaging-based meta-features. By linking the meta-features to scRNA-seq cell types, we found that myeloid cells (clusters 2 and 3) are relatively round and small with high esterase activity; endothelial cells are large and less round as expected, and have intermediate esterase activity; and pericytes have intermediate shape, size and intensity (Figure 4D).

**Figure 4.**
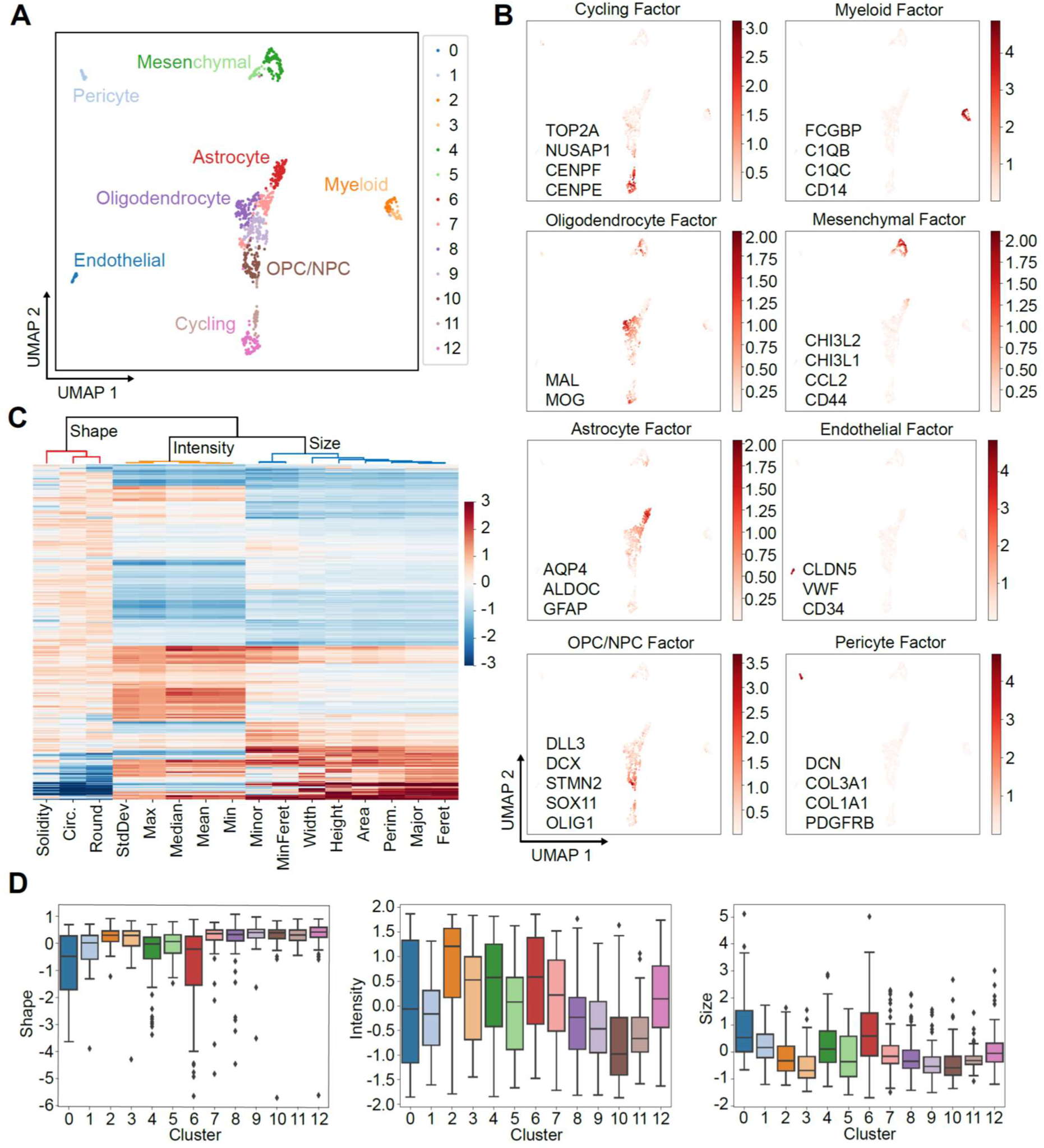
Application of SCOPE-seq2 to a human GBM surgical sample. A) UMAP embedding of the cell scores from scHPF factorization of the scRNA-seq data colored based on unsupervised clustering from Phenograph. B) Same as A) but colored by scHPF cell scores for each scHPF factor. A short list of top-scoring genes for each factor is also included. C) Identification of imaging meta-features. A heatmap shows the z-scored values of 16 cell imaging features (columns) across cells (rows), and a dendrogram indicates three feature clusters, cell size, shape and Calcein staining intensity, from an unsupervised hierarchical clustering. D) Heterogeneity of cell imaging meta-features. Boxplots show the distribution of imaging meta-features in each Phenograph cluster from scRNA-seq.

### SCOPE-seq2 identifies relationships between imaging features and lineage identities of malignantly transformed GBM cells

Malignant cells in GBM can resemble multiple neural/glial lineages or exhibit a mesenchymal phenotype (Neftel et al., 2019; Patel et al., 2014; Verhaak et al., 2010; Yuan et al., 2018a). Because malignant GBM cells are known to be highly plastic, we decided to use a diffusion map to visualize their lineage relationships (Haghverdi et al., 2015). We selected malignant cells based on aneuploidy as described above (Figure 5A), reduced the dimensionality of malignant cell gene expression by scHPF, and visualized the factorized data with a diffusion map, which revealed two major branches (Figure 5B). One branch consists of astrocyte-like cells and terminates with mesenchymal-like cells, while the other branch consists of OPC/NPC cells and cycling cells. This is consistent with our previous studies showing that astrocyte-like and mesenchymal glioma cells are significantly more quiescent than OPC-like glioma cells (Yuan et al., 2018a).

**Figure 5.**
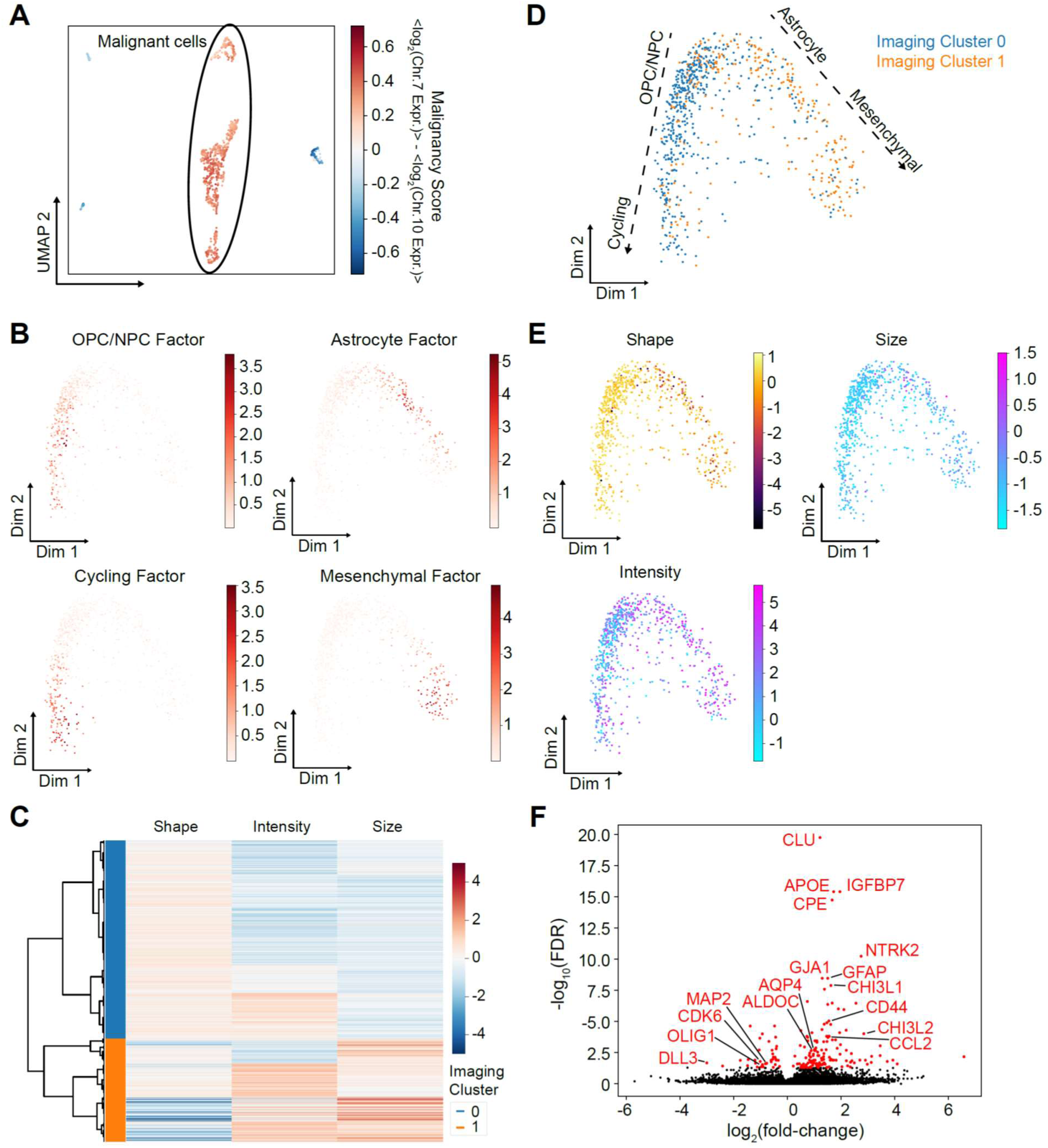
Relationships between cell imaging features and transcriptional phenotypes in GBM. A) UMAP embedding from Figure 4A colored by the malignancy score (the scHPF-imputed difference between Chr.7 and Chr.10 average expression), which indicates malignantly transformed GBM cells based on aneuoploidy. B) Two-dimensional diffusion map of malignantly transformed GBM cells, colored by the scHPF cell scores for factors enriched in GBM lineage markers. C) Clustering of imaging meta-features for the malignantly transformed GBM cells. A heatmap shows the values for three imaging meta-features, and a dendrogram shows the unsupervised hierarchical clustering of cells. Two major imaging clusters of cells are colored. D) Diffusion map in B) colored by the imaging clusters identified in C). E) Diffusion map in B) colored by the values of the three imaging meta-features shown in C). F) Volcano plot for differential expression analysis comparing the two major imaging clusters. Genes with an adjusted p-value (FDR) < 0.05 are indicated in red and many correspond to key markers of the two major GBM branches that were identified.

To explore that how imaging features of malignant cells are related to their transcriptional phenotypes, we asked whether unsupervised clustering of cellular imaging features would correspond to the two major branches observed in the diffusion map from scRNA-seq (Figure 5B). We clustered malignant cells by the three imaging meta-features described above using hierarchical clustering, and identified two major cellular imaging clusters (Figure 5C). By plotting two imaging clusters on the diffusion map embedding of the malignant cells, we found that cells with round shape, low intensity and small size (imaging cluster 0) are enriched in the OPC/NPC-cycling branch, and cells with rough shape, high intensity and large size (imaging cluster 1) are enriched in the astrocyte-mesenchymal branch (Figure 5D, 5E). This finding was further supported by differential expression analysis comparing expression profiles of cells in the two imaging clusters. As expected, markers of OPC/NPCs (MAP2, OLIG1, DLL3) and cycling cells (CDK6) are significantly enriched (FDR<0.05, Mann–Whitney U-test) in imaging cluster 0, while markers of astrocyte-like cells (APOE, GFAP, GJA1, AQP4, ALDOC) and mesenchymal cells (CHI3L1, CD44, CHI3L2, CCL2) are significantly enriched (FDR<0.05, Mann–Whitney U-test) in imaging cluster 1 (Figure 5F, Table EV4). Therefore, there is a clear correspondence between the major gene expression and basic imaging features for the malignantly transformed cells in this tumor.

## Discussion

SCOPE-seq enables a direct link between live cell imaging, cytometry, and scRNA-seq with the scalability and low cost of conventional microfluidic and pooled barcoding approaches. With the upgraded SCOPE-seq2, we achieve enhancements in almost every aspect of performance. In the first experiment described here, we profiled >9,000 cells in a single device, linking >4,100 of them to imaging data. This represents a ∼4-fold improvement in throughput. We also achieved a ∼3-fold improvement in molecular capture efficiency for scRNA-seq and a ∼5-fold decrease in the error rate for linking imaging and sequence data. Importantly, our improved optical barcode design greatly simplified the automation and microscopy required for SCOPE-seq2. In the original SCOPE-seq, a small fraction of the oligonucleotides on each bead contained optical barcodes, which limited the fluorescence intensity of the beads and required the use of laser-based optics, a relatively sensitive camera, and a small field-of-view. Because of the relatively low signal and autofluorescence of the beads, we were restricted to red fluorophores and could only image in a single channel, which limited our multiplexing capacity and decoding speed. In SCOPE-seq2, every oligonucleotide on the bead contains an optical barcode (∼100-fold more oligonucleotides) and so a fast, automated microscope with a large field-of-view camera and simple, LED illumination are sufficient and allow two-color optical decoding. These advances make the technology more accessible and contribute to the improved performance.

SCOPE-seq2 compares favorably to alternative approaches for linking imaging or cytometric data to scRNA-seq. Some of the earliest techniques combined index sorting by FACS with scRNA-seq to link cytometric data with expression profiles on a cell-by-cell basis (Shalek et al., 2013). However, this is relatively expensive, does not allow imaging, and is limited by the scalability of library construction in 96- or 384-well plates. The Fluidigm C1, an early commercial platform for scRNA-seq, which could initially profile tens of cells and later scaled to hundreds of cells, directly linked live cell imaging and scRNA-seq (Lane et al., 2017). However, it was also limited by relatively high operating costs and other performance issues such as multiplet capture (Macosko et al., 2015). The icell8 from Takara/Wafergen can link low resolution imaging for cytometry to scRNA-seq, but throughput is limited to ∼1,000-1,800 cells per sample according to product literature. An upgraded version (Hochgerner et al., 2017) combines the Wafergen technology with FACS and achieves higher throughput (∼7,500 cells per sample), but the imaging resolution appears limited to cytometric applications, likely because the chamber volumes are >1,000-fold larger than in SCOPE-seq2 (hundreds of nanoliters vs. ∼100 picoliters). In addition, achieving a high loading density requires FACS, which can be problematic for some cell types. Relatedly, costs per cell were cited as ∼$1 including 100,000 reads, whereas SCOPE-seq2 is <$0.40 at 100,000 reads due to substantially reduced library preparation costs. Finally, Zhang et al recently reported microfluidic technology for linking cytometric analysis with scRNA-seq using a combination of droplet microfluidics and microfabricated chambers (Zhang et al., 2020). The authors claim that their approach is more scalable than SCOPE-seq, but only demonstrated a throughput of ∼1,200 cells which is fewer than both the initial report of SCOPE-seq and the upgraded version described here. Furthermore, it is unclear whether this approach is able to link live cell imaging with scRNA-seq as opposed to just flow cytometric data.

The improvements to SCOPE-seq2 have enabled applications in primary cells isolated from complex tissues, which are typically more challenging to profile by scRNA-seq than cell lines grown in culture. In parallel with this study, we used SCOPE-seq2 to reveal cell size dynamics during adult neurogenesis in the mouse brain and identify the precise stage in neuronal differentiation where morphological changes associated with cell cycle entry occur(Mizrak et al., 2020). In the same experiment, we also used SCOPE-seq2 to identify the cellular targets of NOTUM, an extracellular WNT antagonist that plays a crucial regulatory role in adult neurogenesis. Here, we profiled a human GBM surgical specimen using SCOPE-seq2. The scRNA-seq data from this experiment recapitulated all of the major cellular populations and states that have been associated with GBM in previous reports. In a focused analysis of the malignantly transformed tumor cells, we discovered a strong correlation between certain morphological features of individual cells and their cellular identities. Consistent with earlier studies, the transformed cells appear to differentiate along two major branches – one that includes OPC-like, NPC-like and proliferative glioma cells, and a second that includes astrocyte-like and mesenchymal-like glioma cells that are more quiescent. Interestingly, these two major branches are also distinguishable by the imaging features of the dissociated cells. The astrocyte-/mesenchymal-like cells are larger, less round, and exhibit higher esterase activity. Differential expression analysis based on imaging classification alone separated canonical markers of these two populations, demonstrating that significant information about cellular identity is encoded in simple imaging observables. To summarize, SCOPE-seq2 is a versatile and high-performance technology for directly linking live cell imaging and scRNA-seq that scales to thousands of cells per sample and enables applications in both cell lines and primary cells dissociated from complex tissues.

## Methods and protocols

### Bead construction

8-nt cell barcode sequences (Table EV1) were designed using an R package ‘DNAbarcodes’ with following criteria: sequences were at least 3 Levenshtein distance from each other; sequences that contain homopolymers longer than 2 nucleotides, with GC content <40% or >60%, or perfectly self-complementary sequences were removed. Full-length mRNA capture oligo sequences (Figure 1B) are then generated with these candidate 8-nt cell barcode sequences in a combinatorial fashion. Self-complimentary score of each candidate 8-nt cell barcode sequence, defined as the length of the longest continuous stretch of self-complimentary sequence among all full-length mRNA capture oligo sequences that contain this 8-nt cell barcode sequence, is computed. Every A-T paring and C-G paring is scored with a length of 2/3 and 1, respectively, to account for the stronger binding affinity of C-G paring compared to A-T paring. The 8-nt cell barcode sequences with the bottom 50% self-complimentary scores are selected.

Bead synthesis was performed by Chemgenes Corp (Wilmington, MA). Toyopearl HW-65S resin (∼30 micron mean particle diameter) (Tosoh Biosciences, cat# 19815, Tosoh Bioscience) with a flexible-chain linker was used as a solid support for reverse-direction phosphoramidite synthesis. Beads were synthesized with sequence ‘TTTTTTTAAGCAGTGGTATCAACGCAGAGTACNN’ at 50 micromole scale, split into 96 parts to add one of the S cell barcode sequences, pooled together to add ‘NN’, split into 96 parts to add one of the Q cell barcode sequences, and pooled together to add ‘NNNN’ and 30 T’s.

### Labeling and Generation of Optical Decoding Probe Pools

192 oligonucleotides that are complementary to the 8-nt cell barcodes (Table EV2) with 3’-amino modifications were synthesized and purified (Sigma-Aldrich), then resuspended in water at 200 μM. To generate probe mixtures corresponding to each bit in the binary code, oligonucleotides labeled with ‘1’ were taken (Figure 1D, Table EV3), pooled and resuspended in 0.1 M sodium tetraborate (pH 8.5) coupling buffer at a final concentration of 22 μM with 0.6 μg/μL reactive fluorophore. Sulfo-CY5 NHS ester (Lumiprobe, cat# 21320) was coupled with S oligo pools, and Sulfo-CY3 NHS ester (Lumiprobe, cat# 23320) was coupled with Q oligo pools overnight at room temperature. Excess fluorophores were removed and oligos were recovered by ethanol precipitation (80% Ethanol, 0.06 M NaCl, 6 μg/mL glycogen). The concentration of probes was quantified using a NanoDrop (Thermo Scientific). Probe pools were diluted such that each probe had a final concentration of ∼20 nM, and the two, distinctly labeled probe pools were mixed together for each binary code bit prior to use.

### SCOPE-seq2

- Preparation
  - A microwell array device was filled with wash buffer (20 mM Tris-HCl pH7.9, 50 mM NaCl, 0.1% Tween-20) and stored in a humid chamber one day before use.
  - Cell culture or tissue samples were dissociated into single cell suspension (see section, GBM tissue processing), and stained with desired fluorescent dyes.
- Cell loading
  - The pre-filled microwell array device was flushed with Tris-buffered saline (TBS).
  - The single cell suspension was pipetted into the microwell array device.
  - After 3-minute, un-trapped cells were then flushed out with TBS.
- Cellular imaging
  - The cell-loaded microwell device was scanned using an automated fluorescence microscope (Nikon, Eclipse Ti2) under the bright-field and fluorescence channels. Bright-field images were taken using an RGB light source (Lumencor, Lida) and wide-field 10x 0.3 NA objective (Nikon, cat# MRH00101). Fluorescence images were taken using LED light source (Lumencor, spectra x), Quad band filter set (Chroma, cat# 89402), wide-field 10x 0.3 NA objective (Nikon, cat# MRH00101) with 470 nm (GFP channel) and 555 nm (TRITC channel) excitation for Calcein AM and Calcein red-orange, respectively.
- scRNA-seq (steps performed on microwell device)
  - Beads (Chemgenes) were pipetted into the microwell device, and untrapped beads were flushed out with 1x TBS. The microwell device containing the cells and the beads was connected to the computer-controlled reagent and temperature delivery system as previously described (Yuan and Sims, 2016).
  - Lysis buffer (1% 2-Mercaptoethanol (Fisher Scientific, cat# BP176-100), 99% Buffer TCL (Qiagen, cat# 1031576) and perfluorinated oil (Sigma-Aldrich, cat# F3556-25ML) was flowed into the device followed by an incubation at 50°C for 20 minutes to promote cell lysis, and then at 25°C for 90 minutes for mRNA capture. Wash buffer supplemented with RNase inhibitor (0.02 U/µL SUPERaseIN (Thermo Fisher Scientific, cat# AM2696) in wash buffer) was flushed through the device to unseal the microwells and remove any uncaptured mRNA molecules.
  - Reverse transcription mixture (1X Maxima RT buffer, 1 mM dNTPs, 1 U/µL SUPERaseIN, 2.5 µM template switch oligo, 10 U/µL Maxima H Minus reverse transcriptase (Thermo Fisher Scientific, cat# EP0752), 0.1% Tween-20) was flowed into the device followed by an incubation at 25°C for 30 minutes and then at 42°C for 90 minutes. Wash buffer supplemented with RNase inhibitor was flushed through the device. The device was disconnected from the automated reagent delivery system.
  - Exonuclease I reaction mixture (1X Exo-I buffer, 1 U/µL Exo-I (New England Biolabs, cat# M0293L)) was pipetted into the device followed by an incubation at 37°C for 45 minutes. TE/TW buffer (10 mM Tris pH 8.0, 1 mM EDTA, 0.01% Tween-20) was flushed through the device.
- Bead optical demultiplexing
  - The microwell device containing the beads with cDNAs was connected to a computer-controlled reagent delivery and scanning system (see section, automated reagent delivery system).
  - Melting buffer (150 mM NaOH) was infused into the device and incubated for 10 minutes. The device was then washed with imaging buffer (2xSSC, 0.1% Tween-20). An automated imaging program scanned the device in the bright-field, Cy3 and Cy5 emission channels. Fluorescence images were acquired using an LED light source (Lumencor, spectra x), Quad band filter set (Chroma, cat# 89402), wide-field 10x objective (Nikon, cat# MRH00101) and 555 nm and 649 nm excitation for Cy3 and Cy5, respectively. Hybridization solution (imaging buffer supplemented with probe pool A, described below) was infused into the device and incubated for 10 minutes. The device was then washed with imaging buffer. An automated imaging program scanned the device in the bright-field, Cy3 and Cy5 emission channels.
  - Repeat the previous step 7 times, with probe pool B to H.
  - Melting buffer was infused into the device and incubated for 10 minutes. The device was then washed with imaging buffer, and then disconnected from the automated reagent delivery system.
- scRNA-seq (steps performed off microwell device)
  - Perfluorinated oil was pipetted into the device to seal the microwells. The device was then cut into 10 regions. Beads from each region were extracted separated by soaking each small piece of bead-containing PDMS in 100% ethanol, vortexing, water bath sonication, and centrifugation in a 1.7 mL microcentrifuge tube. PDMS was then removed by tweezer.
  - Beads extracted from each region were processed in separate reactions for the downstream library construction. Beads were washed sequentially with TE/SDS buffer (10 mM Tris-HCl, 1 mM EDTA, 0.5% SDS), TE/TW buffer, and nuclease-free water. cDNA amplification was performed in 50 µL PCR solution (1X Hifi Hot Start Ready mix (Kapa Biosystems, cat# KK2601), 1 µM SMRTpcr primer (Table EV5)), with 14 amplification cycles (95°C 3 min, 4 cycles of (98°C 20s, 65°C 45 s, 72°C 3 min), 10 cycles of (98°C 20s, 67°C 20s, 72°C 3min), 72°C 5 min) on a thermocycler. PCR product from each piece was pooled and purified using SPRI paramagnetic bead (Beckman, cat# A63881) with a bead-to-sample volume ratio of 0.6:1.
  - Purified cDNAs were then tagmented and amplified using the Nextera kit for *in vitro* transposition (Illumina, FC-131-1024). 0.8 ng cDNA was used as input per reaction. A unique i7 index primer was used to barcode the libraries obtained from each piece of the device. The i5 index primer was replaced by a universal P5 primer (Table EV5) for the selective amplification of 5’ end of cDNA (corresponding to the 3’ end of mRNA). Two rounds of SPRI paramagnetic bead-based purification with a bead-to-sample volume ratio of 0.6:1 and 1:1 were performed sequentially on the Nextera PCR product to obtain sequencing-ready libraries.
  - The resulting single-cell RNA-Seq libraries were pooled and 20% PhiX library (Illumina, FC-131-1024) was spiked-in before sequencing on an Illumina NextSeq 500 with a 26-cycle read 1, 58 cycle read 2, and 8 cycle index read. A custom sequencing primer (Table EV5) was used for read 1.

### Automated reagent delivery system

An automated reagent delivery and scanning system is designed for automated optical decoding. In this system, fixed positive pressure (∼1 psi) stabilized by a pressure regular (SMC Pneumatics, cat# AW20-N02-Z-A) is used to drive fluid flow. The microwell device is constantly pressurized during incubation steps, which prevents evaporation and bubble formation. Two 10-channel rotary selector valves (IDEX Health & Science, cat# MLP778-605) are connected in parallel to toggle between 14 reagent channels. A three-way solenoid valve (Cole-Parmer, cat# EW-01540-11), located at the downstream of the microwell device, is used as an on/off switch for reagent flow. The multi-channel selector valves are controlled by a USB digital I/O device (National Instruments, cat# SCB-68A). The three-way solenoid valve is controlled by the same USB digital I/O device, but through a homemade transistor-switch circuit. The system is controlled by imaging software (Nikon, NIS-Elements).

### Bead optical decoding analysis

Eight cycles of probe hybridizations (A to H) were used for cell barcode optical decoding. For each cycle, the device was imaged in the bright-field, Cy3 and Cy5 emission channels. Beads were first identified in the bright-field image by the ImageJ Particle Analyzer plugin, and the positions of the beads in the bright-field image were recorded. Then the average fluorescence intensities of each bead in the Cy3 and Cy5 images were measured. Beads identified in cycles B to H were mapped to the nearest bead in cycle A. Thus, we obtained a probe hybridization matrix with n beads x 16 intensity values (8 for Cy3 and 8 for Cy5). To call cell barcodes from the imaging data, we tested two methods:

- Cycle-by-cycle

The cycle-by-cycle method was modified from the stage-by-stage decoding method (Gunderson et al., 2004)

- For each cycle and each fluorescent channel;
- Get N log transformed average intensity values;
- Compute an intensity histogram using 50 bins;
- Determine the median intensity value *M*, and identify the highest bin with intensity values smaller than *M* as *B*_1_ and the highest bin with intensity values greater than *M* as *B*_2_;
- Identify the lowest bin *B*_3_ with intensity values between *B*_1_ and *B*_2_;
- Get the medium intensity value *I* of bin *B*_3_, then assign 0 to intensity values smaller than *I* and assign 1 to intensity values greater than *I*.
- Refer to the binary code table. If the code assigned is in the table, then return the corresponding cell barcode sequence.

- Bead-by-bead

The bead-by-bead method was modified from the core-by-core decoding method (Gunderson et al., 2004)

- For each bead and each fluorescence channel;
- Get eight average fluorescence intensity values *x*_1_, *x*_2_, …, *x*_8_;
- Let *y*_1_, *y*_2_, …, *y*_8_ be the sorted values;
- Let *f*_*n*_ = (*y*_*n*+1_ − *y*_*n*_)/*y*_*n*_, *n* = 1, 2, …, 7 be the relative intensity fold change between neighbor sorted values;
- Determine the largest fold change 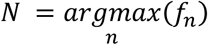, then assign 0 to values to *y*_1_, *y*_2_, …, *y*_*N*_ and assign 1 to values *y*_*N*+1_, *y*_*N*+2_, …, *y*_8_;
- Refer to the binary code table. If the code assigned in step 4 is in the table, then return the corresponding cell barcode sequence;
- Otherwise, remove *f*_*N*_ from list {*f*_*n*_} and repeat step 4, 5, until a corresponding cell barcode sequence is returned or the list {*f*_*n*_} is empty.

### Live cell imaging analysis

Images were analyzed using the ImageJ software as previously described (Yuan et al., 2018b).

- Identify microwells with cells Microwell outlines were identified as objects from the bright-field image using a local threshold, and then average fluorescence intensities of microwells in the live staining images were measured. Average intensity values followed a bimodal distribution, with the higher intensity population corresponding to microwells that contain cells.
- Cell optical phenotype extraction Only microwells with cells were selected and each cell was analyzed individually within the smallest bounding square of the corresponding microwell. The cell was identified in the live staining fluorescence image using the auto threshold and particle analyzer. Microwells with multiple cells identified by the software were excluded. Sixteen imaging features were measured for each cell in the fluorescence image: area, mean intensity, standard deviation of intensity, minimum intensity, maximum intensity, median intensity, perimeter, width, height, major axis, minor axis, circularity, Feret’s diameter, minimum Feret’s diameter, roundness, and solidity.

### SCOPE-seq2 scRNA-seq analysis

To analyze the scRNA-seq data from SCOPE-seq2, we first extracted the cell-identifying barcode and UMI from Read 1 based on the designed oligonucleotide sequence, NN(8-nt Cell Barcode S)NN(8-nt Cell Barcode Q)NNNN. The 192 8-nt cell barcode sequences have a Hamming distance of at least three for all sequence pairs. Therefore, we corrected one substitution error in the cell barcode sequences. We only keep reads with a complete cell barcode. Next, we align the reads from Read 2 to a merged human/mouse genome (GRCh38 for human and GRCm38 for mouse) with merged GENCODE transcriptome annotations (GENCODE v.24 for both species) using STAR v.2.7.0 aligner (Dobin et al., 2013) after removing 3’ poly(A) tails (indicated by tracts of >7 A’s) and fragments with fewer than 24 nucleotides after poly(A) tail removal. Only reads that were uniquely mapped to exons on the annotated strand were included for the downstream analysis. Reads with the same cell barcode, UMI (after one substitution error correction) and gene mapping were considered to originate from the same cDNA molecule and collapsed. Finally, we used this information to generate a molecular count matrix.

### SCOPE-seq2 linking cell imaging and sequencing data

To link cell barcodes identified from imaging to cell imaging phenotypes, bright-field images of the device obtained during optical decoding were mapped to images of the live cell imaging based on the upper-left and the bottom right microwells. Cells were then registered to the nearest mapped bead within a microwell radius. To link cell imaging phenotypes to expression profiles, we only considered cell barcodes with registered cells, then we found the exact and unique mapping of the cell barcodes from imaging and sequencing.

### Cell culture

Human U87 and mouse 3T3 cells are cultured in Dulbecco’s modified eagle medium (DMEM, Life Technologies, cat# 11965118) supplemented with 10% fetal bovine serum (FBS, Life Technologies, cat# 16000044) at 37°C and 5% carbon dioxide.

### GBM tissue processing

A single-cell suspension was obtained from excess material collected during surgical resection of a WHO Grade IV GBM. The patient was anonymous and the specimen was de-identified. The tissue was mechanically dissociated following a 30-minute incubation with papain at 37°C in Hank’s balanced salt solution. Cells were re-suspended in TBS after centrifugation at 100xg followed by selective lysis of red blood cells with ammonium chloride for 15 minutes at room temperature. Finally, cells were washed with TBS and quantified using a Countess (ThermoFisher).

### Human and mouse cells mixed experiment

- Human U87 cells were stained with Calcein AM (ThermoFisher Scientific, cat# C3100MP) and mouse 3T3 cells were stained with Calcein red-orange (ThermoFisher Scientific, cat# C34851) in culture medium at 37°C for 10 minutes. The stained cells are then dissociated into single cell suspension by 0.25% Trypsin-EDTA (Life Technologies, cat# 25200-072) and re-suspended in TBS buffer. The U87 and 3T3 cells were mixed at 1:1 ratio with a final total cell concentration 1000 cells/μl.
- The mixed cell suspension was processed and sequenced with SCOPE-seq2 workflow described above (PJ070).
- Images and sequencing data were processed with the SCOPE-seq2 pipeline described above.

### Sub-sampling analysis

To analyze the saturation behavior and sensitivity of scRNA-seq data from SCOPE-seq2 (Figure 3A), we randomly sub-sampled the aligned reads and re-processed them with the scRNA-seq analysis pipeline described above. We then calculated two statistics, molecules per cell and genes per cell, based on the cells that were discovered from the total reads.

### Accuracy of linking imaging and scRNA-seq data

The linking accuracy was defined as the concordance between the scRNA-seq and imaging-based species calling for cell barcodes associated with a single species. In scRNA-seq data, cells with >90% of reads aligning uniquely to a given species were considered to correspond to a single species. In the imaging data, we determined the imaging-based species call based on cell live staining colors. Cells with Calcein AM intensity > 724 were called as imaging-based human cells; Cells with Calcein red-orange intensity > 2,048 were called as imaging-based mouse cells. Intensity thresholds were determined as the intensity of the shortest bin between the two mean values of the bimodal Gaussian distribution of intensity values.

### Imaging based multiplets identification

Two-color live staining fluorescence images were merged with Calcein AM signal in green and Calcein red-orange signal in magenta. Each well was manually examined within the smallest bounding square. Wells with mixed-species cells were determined as having at least one green object and one magenta object; wells with a single cell were determined as having only one green object or one magenta object.

### Glioblastoma experiment

- GBM specimen was collected and dissociated into single cells as described above. Cells were stained with Calcein AM (ThermoFisher Scientific, cat# C3100MP)
- The GBM cell suspension was processed and sequenced with SCOPE-seq2 workflow described above (PJ069).
- Imaging and sequencing data were processed with the SCOPE-seq2 pipeline described above.
- We removed multiplets based on manually examination of each well within the smallest bounding square of the Calcein AM fluorescence image.
- We identified the dead cells based on the Calcein AM fluorescence intensity. We fitted a Gaussian distribution to the fluorescent intensity histogram, set a threshold of lower 5 percentile, and removed cells with intensity lower than the threshold.

### Single cell hierarchical Poisson factorization (scHPF) analysis

To reduce the dimensionality of scRNA-seq results, we factorized gene count matrix using the scHPF (Levitin et al., 2019) with default parameters and K = 13 (www.github.com/simslab/schpf). One of the factors contained several heat shock with high gene scores (among the top 50 genes), likely indicating dissociation artifacts in certain cells. This factor was removed in all downstream analysis.

### Malignant cell identification

The cell aneuploidy analysis was performed based on the scHPF model as described previously (Zhao et al., 2020). To compute the scHPF-imputed expression matrix, we multiplied the gene and cell weight matrix (expectation matrix of variable θ and β) in the scHPF model and then log-transformed the result matrix as *log*_2_(*expected counts*/10000 + 1). The average gene expression on each somatic chromosome was calculated using the scHPF-imputed count matrix as previously described (Yuan et al., 2018a). We defined a malignancy score as the difference between the average expression of Chr. 7 genes to that of Chr. 10 genes, < *log*_2_(*Chr*. 7 *Expression*) > − < *log*_2_(*Chr*. 10 *Expression*) >, plotted in Figure 5A. We fitted a double Gaussian distribution to the malignancy score and the score of the shortest bin between two mean intensities was used as the threshold that separates the malignant and non-malignant cell populations (Figure EV3A). The difference of chromosome average expression between malignant and non-malignant cells, computed as the expression subtracted by the average expression of non-malignant cells, was shown in Figure EV3B.

### scRNA-seq clustering and visualization

To visualize the scHPF model (Figure 4A), we generated a UMAP embedding using the Pearson correlation distance matrix computed from the cell score matrix. To cluster the scRNA-seq profiles, we used the Phenograph implementation of Louvain community detection (Levine et al., 2015), with the Pearson correlation matrix and k=50 to construct a k-nearest neighbors graph.

### Cell optical phenotypes clustering

To reduce the dimensionality of the cellular imaging features, 16 cell imaging features were z-normalized and hierarchically clustered using the ‘linkage’ method in the python module ‘scipy’ with correlation distance. The dendrogram in Figure 4C was cut as k=3 to form three clusters of imaging features, corresponding to cell size, shape and esterase activity. The values of meta-features were calculated as an average of the imaging features within each cluster.

To cluster the malignant cells based on their optical phenotypes, we hierarchically clustered imaging meta-features using the ‘linkage’ method in the python module ‘scipy’ with correlation distance. The dendrogram in Figure 5C was cut as k=2 to form two clusters of malignant cells.

### Diffusion map embedding of malignantly transformed GBM cells

We factorized the molecular count matrix for malignantly transformed GBM cells (identified by aneuploidy analysis as described above) using scHPF with default parameters and K=15. Prior to further analysis, we removed one of the 15 factors, which exhibited high scores for heat shock response genes, because it likely represents a dissociation artifact in a subset of cells. We then computed diffusion components with the DMAPS Python library (https://github.com/hsidky/dmaps). A Pearson correlation distance matrix computed from the scHPF cell score matrix was used as input with a kernel bandwidth of 0.5. The first two diffusion components are plotted in Figure 5B,D,E.

### scRNA-seq differential expression

We used the Mann-Whitney U-test for differential expression analysis. For pairwise comparison of two groups of cells, the group with more cells was randomly sub-sampled to the same cell number as the group with fewer cells. Next, the detected molecules from the group with a higher average number of molecules detected per cell were randomly sub-sampled so that the two groups had the same average number of molecules detected per cell. The resulting sub-sampled matrices were then normalized using the random pooling method from Lun et al as implemented in the scran R package (L. Lun et al., 2016). Finally, the resulting normalized matrices were subjected to gene-by-gene differential expression testing using the Mann-Whitney U-test using the ‘mannwhitneyu’ function in the Python package SciPy. The resulting p-values were corrected using the Benjamini-Hochberg method as implemented in the ‘multipletests’ function in the Python package statsmodels.

## Acknowledgements

P.A.S. was funded by R33CA202827 from NIH/NCI, R44HG010003 from NIH/NHGRI, and a Human Cell Atlas Pilot Project grant from the Chan Zuckerberg Initiative. P.A.S., A.L., and A.I. were funded by U54CA193313 from NIH/NCI. P.A.S., P.C., and J.N.B. were funded by R01NS103473 from NIH/NINDS. This research was funded in part through the NIH/NCI Cancer Center Support Grant P30CA013696 and used the Genomics and High Throughput Screening Shared Resource.

## Author contributions

J.Y., Z.L and P.A.S. conceived the study and designed the experiments. J.Y., Z.L. and P.A.S. developed the SCOPE-seq2 experimental protocols. J.N.B., and P.C. procured the GBM tissue. A.L. and A.I. prepared glioma samples. Z.L., J.Y. and P.A.S. analyzed the data. Z.L. and P.A.S. wrote the manuscript. All authors edited, read and approved the final manuscript.

## Data availability

The datasets produced in this study are available in the following database: scRNA-seq data: Gene Expression Omnibus GSE151137 (https://www.ncbi.nlm.nih.gov/geo/query/acc.cgi?acc=GSE151137). Code is available at: https://github.com/simslab.

