## Supplementary Figures and Table Legends for "Integrating single-cell RNA-seq and imaging with SCOPE-seq2"

Table EV1. Cell barcode sequences.

Table EV2. Fluorescent probe sequences for cell barcode optical decoding.

Table EV3. Cell barcode temporal binary codes.

Table EV4. Differential expression of two imaging clusters.

Table EV5. Primer sequences.

### Source Data Legends

Source Data for Figure 4. Gene score matrix from the scHPF model of all GBM cells.

Source Data for Figure 4C. Z-scored cell imaging features.

Source Data for Figure 5. Gene score matrix from the scHPF model of malignantly transformed GBM cells.

### Extended View Figures

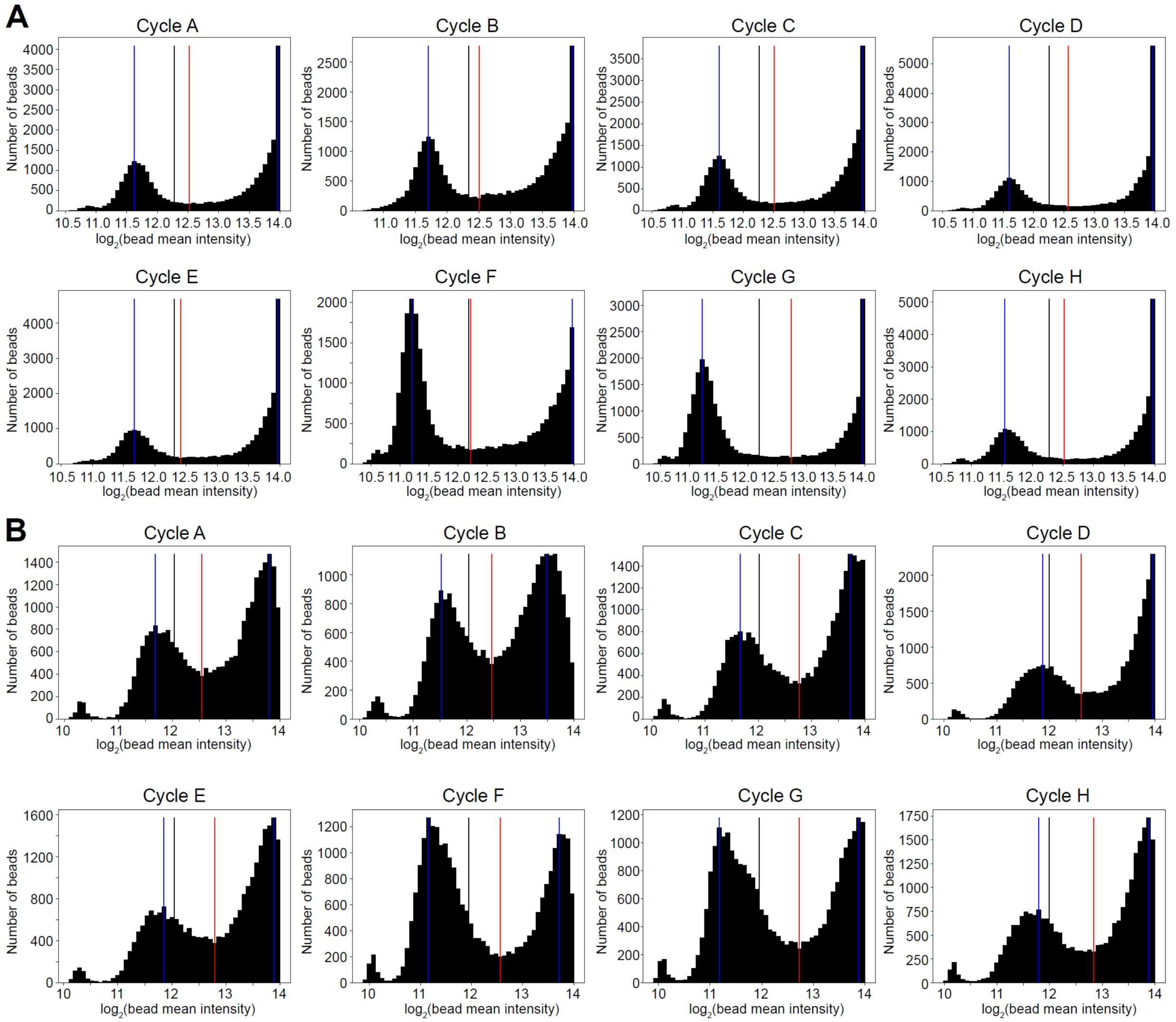


**Figure EV1 - Cell barcode optical decoding.** A), B) Histograms show fluorescence intensity distributions of probe hybridizations for cell barcode S, which are labeled with Cy5 (A) and for cell barcode Q, which are labeled with Cy5 (B) in data set PJ070. Black vertical line shows the median bin; blue vertical line shows the highest bins on each side of the black vertical line; red vertical line shows the lowest bin, the threshold to separate ‘zero’ and ‘one’ populations, between the two blue vertical lines.

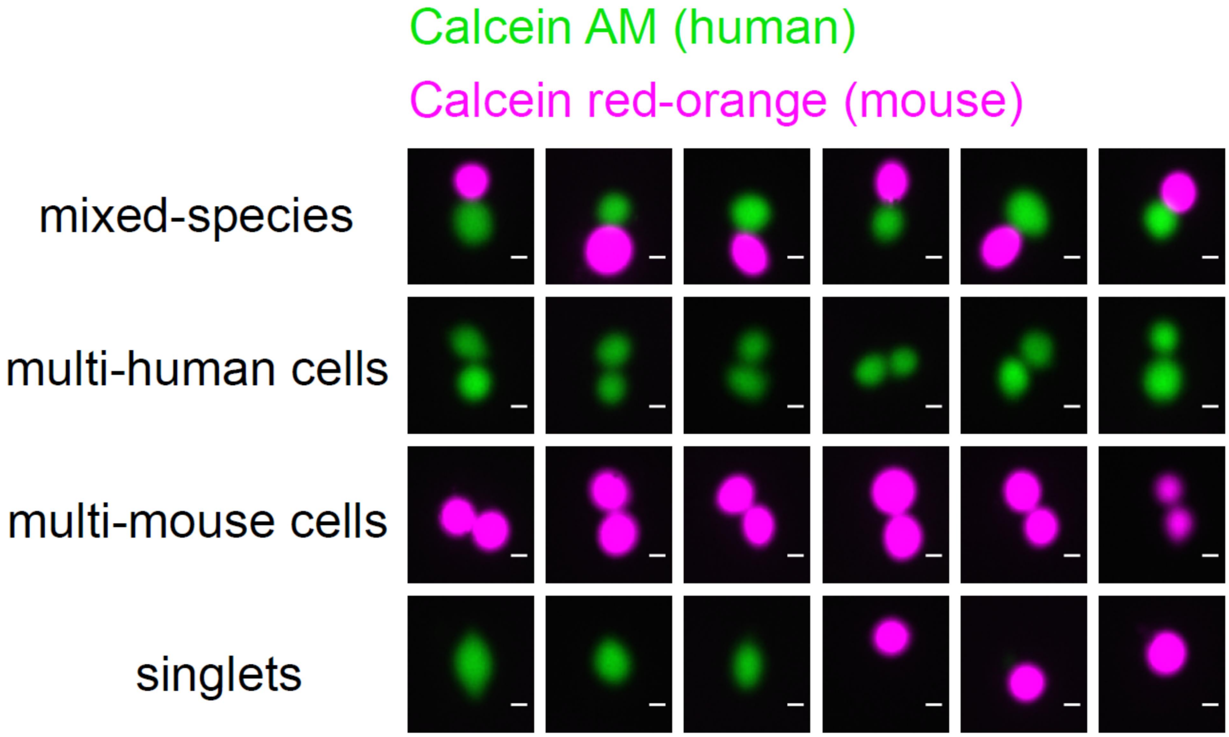


**Figure EV2 - Multiplet detection.** Fluorescence images show example wells with mixed-species cells, multi-human cells, multi-mouse cells and singlets. Scale bar: 10 µm.

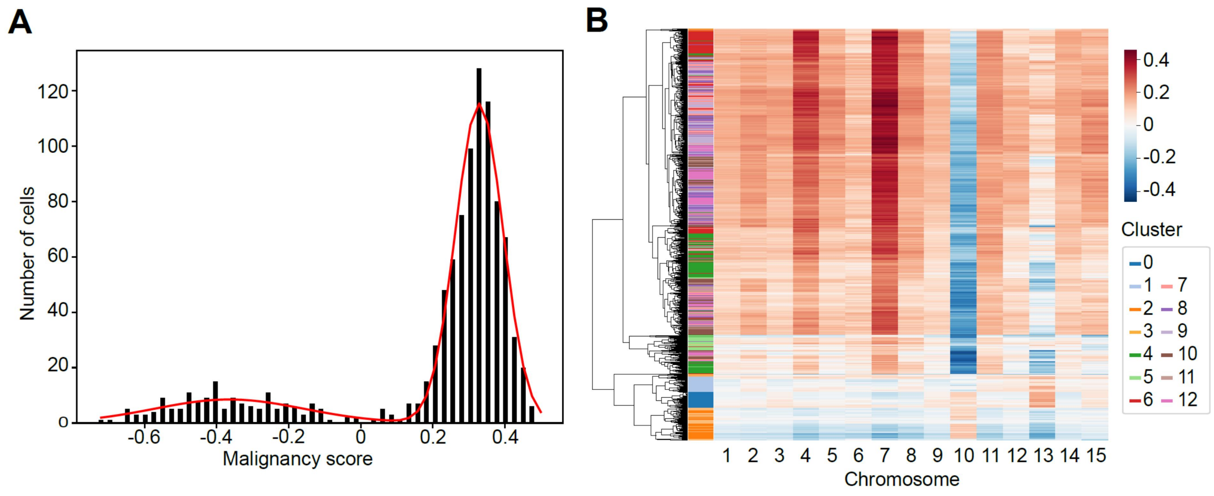


**Figure EV3 – Malignantly transformed GBM cell identification.** A) Histogram shows the distribution of malignancy scores across all cells. B) Heatmap shows the relative chromosomal average expression (chromosomal average expression with subtracted non-malignant cell chromosome average expression subtracted). Colors indicate the Phenograph cell clusters.
